# Efficiency of the synthetic self-splicing RiboJ ribozyme is robust to *cis*- and *trans*-changes in genetic background

**DOI:** 10.1101/2021.03.17.435894

**Authors:** Markéta Vlková, Bhargava Reddy Morampalli, Olin K. Silander

## Abstract

The expanding knowledge of the variety of synthetic genetic elements has enabled the construction of new and more efficient genetic circuits and yielded novel insights into molecular mechanisms. However, context dependence, in which interactions between proximal (*cis*) or distal (*trans*) elements affect the behaviour of these elements, can reduce their general applicability or predictability. Genetic insulators, which mitigate unintended context-dependent *cis*-interactions, have been used to address this issue. One of the most commonly used genetic insulators is a self-splicing ribozyme called RiboJ, which can be used to decouple upstream 5’ UTR in mRNA from downstream sequences (e.g., open reading frames). Despite its general use as an insulator, there has been no systematic study quantifying the efficiency of RiboJ splicing or whether this autocatalytic activity is robust to *trans*- and *cis*-genetic context. Here, we determine the robustness of RiboJ splicing in the genetic context of six widely divergent *E. coli* strains. We also check for possible *cis*-effects by assessing two SNP versions close to the catalytic site of RiboJ. We show that mRNA molecules containing RiboJ are rapidly spliced even during rapid exponential growth and high levels of gene expression, with a mean efficiency of 98%. We also show that neither the *cis*- nor *trans*-genetic context has a significant impact on RiboJ activity, suggesting this element is robust to both *cis*- and *trans*-genetic changes.

## Introduction

Synthetic nucleic acid functional elements used to control protein output - such as promoters, ribosome binding sites, or terminators - are an indispensable part of engineered genetic circuits (Levskaya et al., 2005; Na et al., 2013; Neves et al., 2020), and are frequently used to study basic biological processes (Barbier et al., 2020; Bittihn et al., 2020). The value of such synthetic functional elements increases as their properties are better described and quantified - in some cases, careful quantification of the behaviour of synthetic functional elements has led to fundamentally new insights into molecular mechanisms controlling protein output (Schmiedel et al., 2019; Urtecho et al., 2019).

One important type of synthetic functional element that has been used to ensure predictable and robust protein output from mRNA are self-splicing ribozymes. These ribozymes can be used to splice mRNA at specific locations, for example to remove the 5’ untranslated region (UTR). One of the most common ribozymes used to remove 5’ UTRs is RiboJ. By removing the 5’ UTR of the mRNA, RiboJ enables transcripts having different promoters and thus different 5’ UTRs to produce identical mRNAs. This mitigates any effects that the 5’ UTR might have on mRNA folding or ribosome binding, keeping the translation initiation rate consistent (and predictable) even when promoters have different sequences (Neves et al., 2020; Urtecho et al., 2019; Yu et al., 2018). The utility of the RiboJ element was first demonstrated when it was used to ensure predictable expression in a synthetic NOT gate circuit, irrespective of the sequence of the promoter used to control the expression of a CI repressor in the system (Lou et al., 2012). However, since the first use of RiboJ as a means of ensuring predictable expression, additional research has suggested there can also be unexpected effects of its use. (Clifton et al., 2018) demonstrated that RiboJ insertion into the mRNA sequence led to an increase in protein expression, and that the relative increase in expression depended on the strength of the promoter used. This effect was attributed to hairpin formation at the 5’ end of mRNA whose 5’ UTR had been removed by RiboJ, leading to higher stability and increased translation (Carrier & Keasling, 1997; Clifton et al., 2018; Neves et al., 2020). Another unexpected effect was observed when (Bartoli et al., 2020) designed a tunable system to control translation initiation via binding of small regulatory RNA (sRNA). The complex secondary structure of mRNA molecules with RiboJ at the 5’ end appeared to interfere with the sRNA binding, decreasing the performance of the system. These results emphasise that unknown properties and behaviors of synthetic functional elements - here, RiboJ in particular - can lead to unexpected obstacles when creating new synthetic circuits.

We hypothesised that a complicating factor in the use of the RiboJ system would be the varying efficiency of RiboJ autocatalytic splicing activity between different bacterial strains, or due to polymorphisms near the RiboJ element. To our knowledge, there has been no systematic study quantifying the efficiency of the autocatalytic RiboJ splicing, or whether the efficiency of this autocatalytic activity depends on the genetic background of the organism in which it is used. To address these questions we first developed an assay to quantify RiboJ self-splicing efficiency. We then tested the robustness of the self-splicing activity to *cis*-genetic changes by assaying efficiency in two genetic contexts that differ by a single nucleotide polymorphism (SNP) close to the autocatalytic site of RiboJ. Finally, we tested the robustness of RiboJ behaviour to *trans*-genetic changes by quantifying efficiency in six widely divergent strains of *E. coli*.

Our first motivation for quantifying the behaviour of RiboJ arose during experiments aimed at quantifying the effects of polymorphisms in the *lacZ* promoter on protein expression. We discovered a single SNP at position 69 relative to the *lacZ* gene start codon (C to A) that resulted in a change in downstream protein levels. We quantified this effect by placing the *lacZ* promoter upstream of GFP and measuring fluorescence in a flow cytometer. The effect of this SNP on GFP expression was apparent in different genetic backgrounds as well as during growth in different carbon sources (**Fig. 1A**). To check whether this C69A SNP affected transcription or translation (or both), we incorporated RiboJ as an insulator downstream of it(**Fig. 2**, Top panels). This ensured that the mRNA being translated was identical regardless of which SNP was present. Thus, if the change in protein expression remained in the presence of RiboJ, we could infer that the change was due solely to the SNP affecting transcription. If the difference disappeared in the presence of RiboJ, we could infer that the change was due to the SNP affecting translation. However, it was also possible that the SNP itself interfered with RiboJ cutting (a *cis*-effect). If so, we could not unambiguously infer the cause of the changes in fluorescence we observed were due to translation or transcription (or both). In addition, it is possible that the genetic background of the strain itself affected RiboJ cutting (a *trans*-effect). To exclude the possibility of *cis*- or *rans*-effects on RiboJ cutting efficiency, we quantified efficiency in the presence of *cis*- and *rans*-genetic changes.

**Figure 1:**
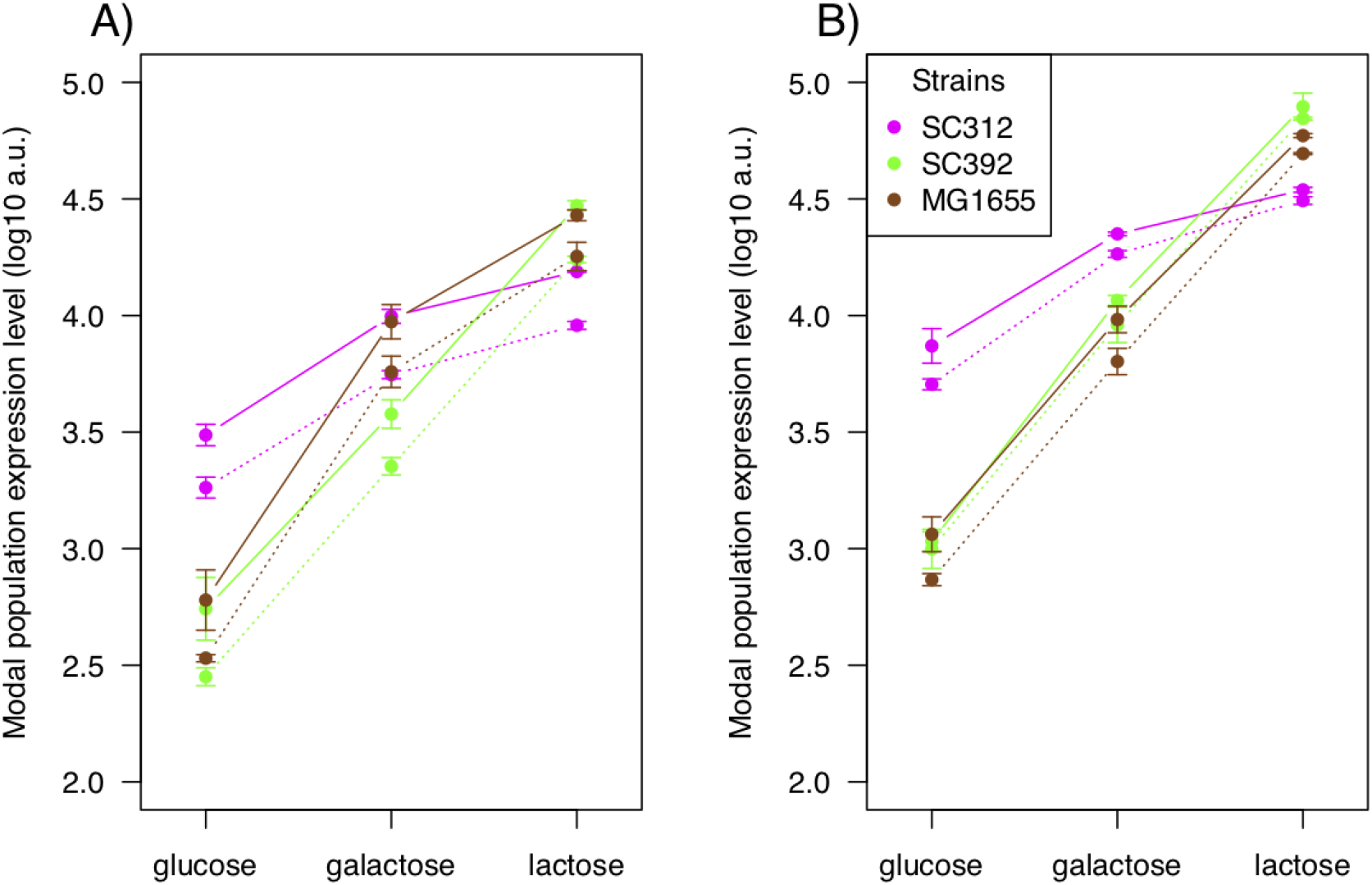
Modal expression levels differ consistently due to a single A to C change at position +69 of the *lacZ* open reading frame both without RiboJ (A) and with RiboJ (B) and across genetic backgrounds. Shown are the modal population fluorescence levels for GFP driven by an upstream *lacZ* promoter for three divergent strains of *E. coli*. In all genetic backgrounds tested and all growth conditions (glucose, galactose, and lactose), the fluorescence levels from a promoter with the 69C polymorphism (dotted line) were consistently lower than those from promoter with the 69A polymorphism (solid line). There are clear effects of genetic background on both the level and dynamic range of protein expression. In particular, SC312 has a narrow dynamic range, with relatively high expression in non-lactose environments compared to the other strains, but relatively low expression in lactose. Despite this, the effect of the A to C change is nearly constant. Whiskers show one standard deviation of three replicates.

**Figure 2:**
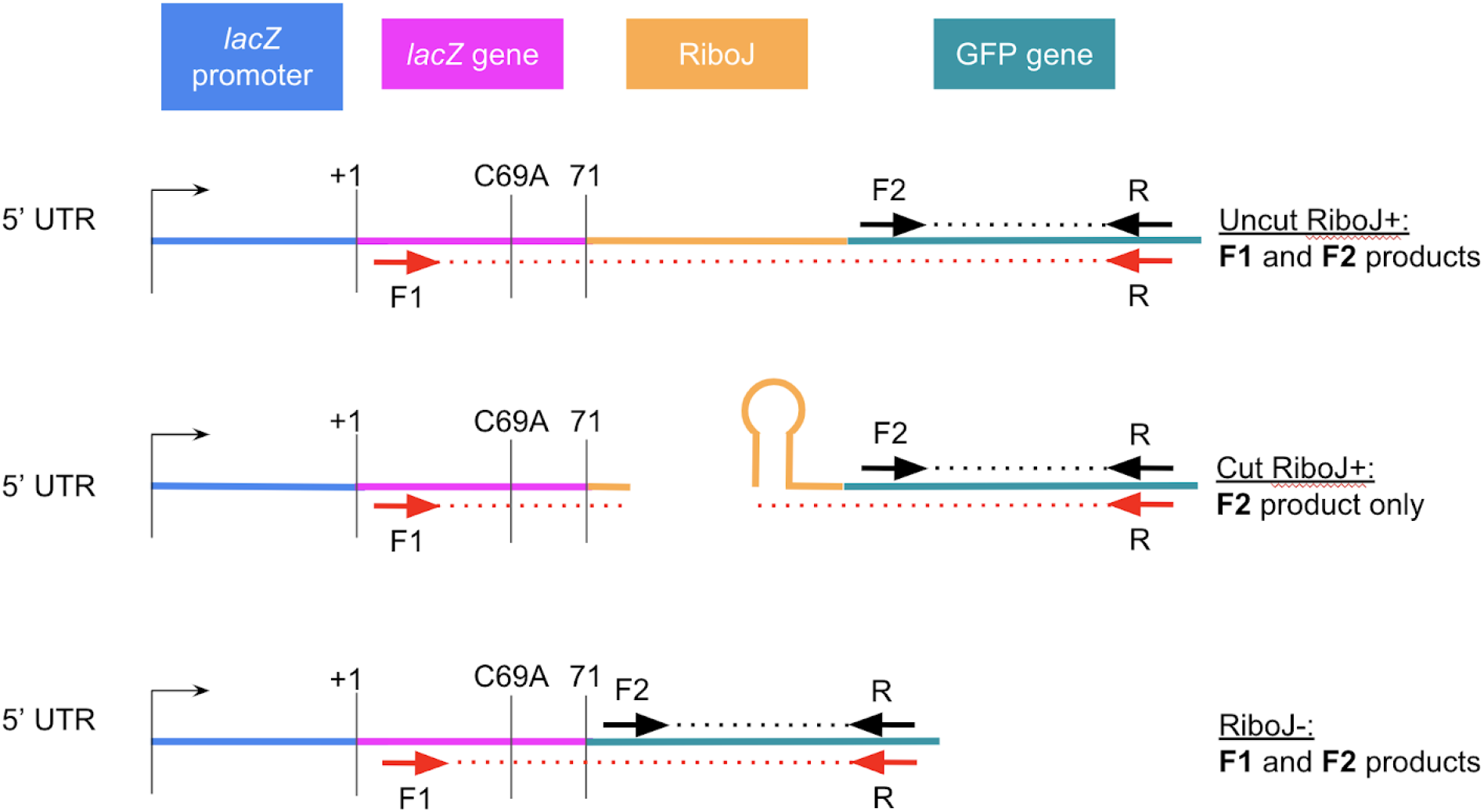
Scheme of the RT-qPCR primer design to quantify the efficiency of RiboJ cutting. Each primer is represented by an arrow, with pairs coloured the same. The dotted lines indicate amplicons. If the dotted line between a primer pair is interrupted, the amplicon is not produced. When RiboJ cleaves off the 5’ UTR (Middle panel), the amplicon from primer F1 is not produced, while the amplicon from F2 primer is still produced. The *lacZ* promoter and first 71bp of the *lacZ* open reading frame were placed upstream of GFP (and RiboJ) as a part of 5’ UTR. The translation is driven from a strong synthetic ribosome binding site downstream of the *lacZ* gene sequence (here as a part of the GFP gene).

We designed an RT-qPCR assay to quantify the autocatalytic cutting activity of RiboJ. This assay is based on the principle that for a pair of forward and reverse primers that span the RiboJ cut site, an amplification product should only be produced for uncut mRNA molecules. In contrast, for primer pairs that do not span the cut site, an amplification product should be produced for all molecules. By examining the relative numbers of cut and uncut molecules, we can infer the efficiency of RiboJ cutting relative to the rate of production of all transcripts controlled by the same promoter (i.e. the rate of transcription). To this end, we designed two qPCR primer sets. The first set produced an amplicon from a region spanning the RiboJ cut site, while the second produced an amplicon from a region downstream of the RiboJ cut site (**Fig. 2**). Both sets shared the same reverse primer, differing solely by the location of forward primer. Because one forward primer binds upstream of the RiboJ cut site, no amplification can occur if the 5’ UTR sequence has been cut off (**Fig. 2**, Middle panel). The second forward primer binds downstream of the cut site and results in an amplification product from all transcripts. To quantify differences in amplification that might result from primer binding or other unforeseen mechanisms, we calculated the relative fold change in the abundance of these two amplicons when RiboJ is absent. In the absence of RiboJ, any difference in amplification between the two primer sets should be due solely to differences in primer efficiency or related effects, as without RiboJ, both amplification products will always be produced (**Fig 2.** Bottom panel).

We first assessed whether *trans*-genetic changes affected the self-splicing activity of RiboJ, by assaying RiboJ activity in six widely divergent strains of *E. coli* (**Table 1**). To test for *cis*-effects, we assayed activity in two promoter contexts, each varying by a single SNP 8bp upstream of the RiboJ cut site (2bp upstream of RiboJ sequence). We thus transformed each of the six strains with each of four plasmids differing by the C69A SNP in the *lacZ* promoter and either with RiboJ or without RiboJ (**Fig. 2**, **Table 2**). We isolated RNA from exponentially growing cultures for all strains and confirmed that the amplification efficiency of all primer combinations with all templates was within the range of 90 - 110% (**Fig. S1**). We used the resulting mean efficiency value across all strains (95.8%) for all subsequent calculations of RiboJ autocatalytic activity. We assayed the efficiency of RiboJ autocatalytic activity using at least triplicates for each strain and promoter combination (**Methods**). RiboJ cutting efficiency was high in all cases. Overall we found that 98% of all mRNA molecules containing RiboJ were cleaved. This was extremely robust for almost all strain and promoter combinations, with the lowest median value being 97% (**Fig. 3**). We also found that RiboJ activity was robust to *cis*-changes, with no consistent differences between the 69A and 69C versions of the promoter.

**Figure 3:**
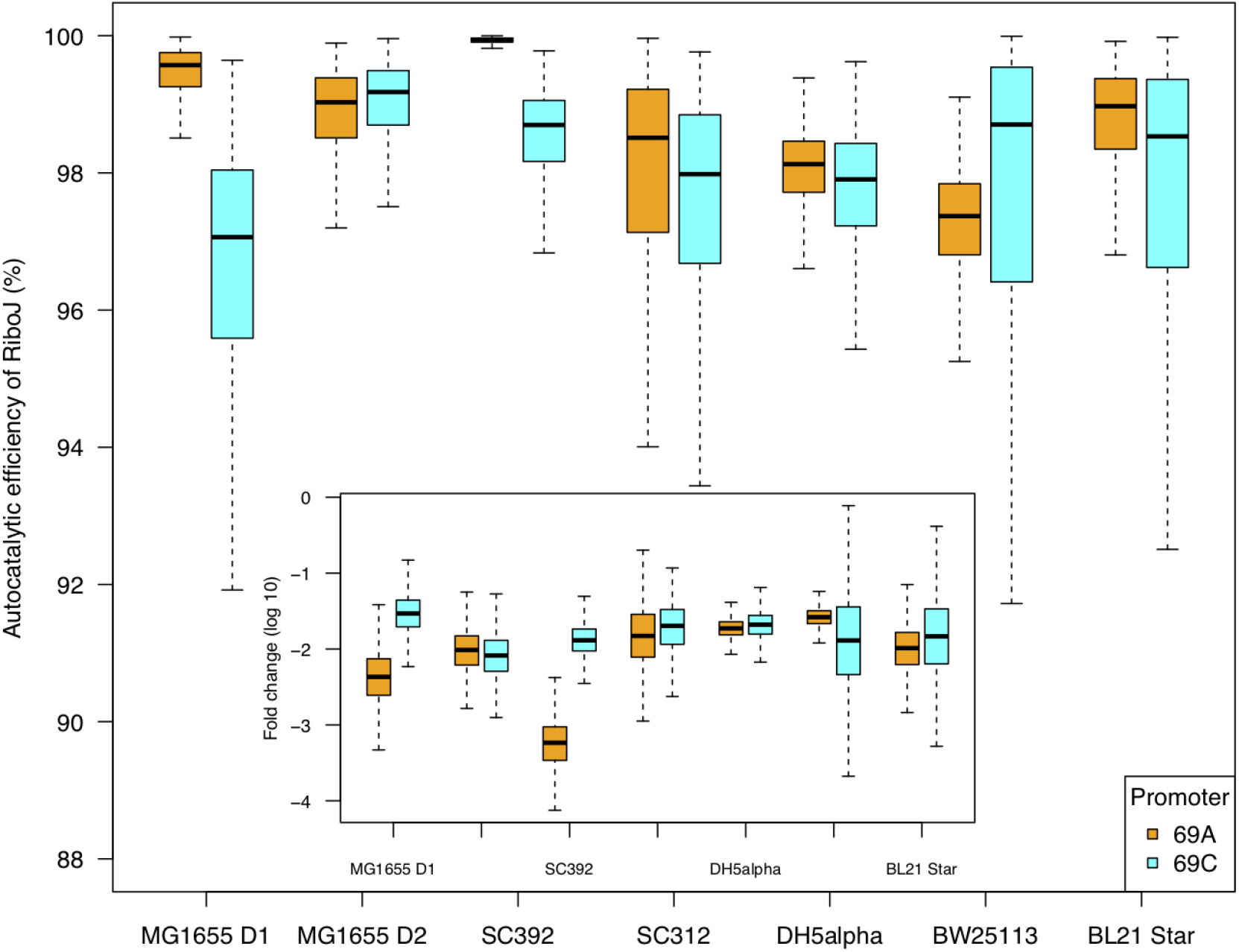
Autocatalytic efficiency of RiboJ. The boxplots in the figure show the minimum and maximum value (whiskers), the first and third quartile (boxes), and the median. These values were obtained through bootstrapping the RT-qPCR data (see Methods). D1 and D2 in MG1655 strain labels indicate that this data is from different biological replicates for which the RNA was extracted on different days (D1 and D2 denoting day 1 and day 2, respectively). The inset shows fold-changes in abundance of uncut transcripts with RiboJ relative to all transcripts. Note that the smaller range in cutting efficiency of RiboJ in SC392 strain for promoter 69A is simply a consequence of converting the C_t_ fold-change of the two different amplicons into catalytic efficiency in percentages. The inset shows that the range and error in fold-changes for SC392 with promoter 69A is comparable to the other samples.

**Table 1:** Bacterial strains and plasmids used in this study. All plasmids carry Kan^R^ selection marker and were created using pUA66 backbone (Zaslaver et al., 2006).

| **Bacterial strains** |  |  |  |
| --- | --- | --- | --- |
| Strain | Relevant characteristics | Phylogroup | Source or reference |
| SC392 | *Natural isolate of E. coli; Soil; 7/18/05; SC15-U2out14; St. Louis Clyde; Upshore (2m) outside the box* | B1 | (Ishii et al., 2006) |
| SC312 | *Natural isolate of E. coli; Water; 6/15/05; SC14-W8; St. Louis Clyde; Surface water* | B1 | (Ishii et al., 2006) |
| MG1655 | *F- λ-ilvG- rfb-50 rph-1* | A | (Blattner, 1997) |
| DH5α | *F- ϕ80lacZΔ M15 $\Delta$ (lacZYA-argF) U169 recA1 endA1 hsdR17 (rK- mK+) phoA supE44 λ-thi-1 gyrA96 relA1* | A | Invitrogen |
| BW25113 | *F- DE(araD-araB)567 lacZ4787(del)::rrnB-3 LAM-rph-1 DE(rhaD-rhaB)568 hsdR514* | A | (Datsenko & Wanner, 2000) |
| BL21 Star (DE3) | *F-ompT hsdSB (rB-, mB-) galdcmrne131 (DE3)* | A | Invitrogen |
| **Plasmids** |  |  |  |
| Plasmid | Relevant characteristics | Source |  |
| p69A.RJ- | *lacZ* promoter 69A, without RiboJ | D. Blank, University of Basel |  |
| p69C.RJ- | *lacZ* promoter 69C, without RiboJ | D. Blank, University of Basel |  |
| p69A.RJ+ | *lacZ* promoter 69A, with RiboJ | This study |  |
| p69C.RJ+ | *lacZ* promoter 69C, with RiboJ | This study |  |

However, we observed one exception to this robust behaviour. In strain SC392, the 69A version of the promoter construct exhibited a 10-fold higher cutting efficiency (**Fig. 3**, inset). To obtain an estimate of error for our measurements and test whether sampling alone could account for this or other differences, we bootstrapped the data 10,000 times and re-calculated the efficiencies (**Fig. 3**, **Methods**). We found that sampling alone was unlikely to account for the higher efficiency of 69A that we observed. This raises the possibility that the *cis*-genetic change only has an observable effect in some genetic backgrounds. However, in light of the differences we observed between MG1655 samples extracted on different days (**Fig. 3**), and the robust behaviour of RiboJ in all other genetic backgrounds, it is also possible that *cis*-effect observed in strain SC392 may be due to some unforeseen artifact or other factor, such as a mutation elsewhere in some of the plasmids.

Finally, the robust behaviour of RiboJ to changes in both *cis*- and *trans*-genetic contexts that we observed, suggests that the differences in expression that we observed in **Fig. 1** are a result of changes in both transcription and translation due to the single C69A SNP, and not to changes in RiboJ autocatalytic efficiency. We note that there have been no previous reports that this region is involved in *lacZ* gene regulation.

## Materials and Methods

### Bacterial strains

The genetic backgrounds of *E. coli* strains used in this study are listed in **Table 1**. The identity of all lab strains was confirmed using whole genome sequencing. The whole genomes of strains SC312 and SC392 have been also sequenced (Breckell & Silander, 2020). Four different plasmids (**Table 1**) were transformed into each of the strains, providing 24 clones that we used to evaluate the efficiency of RiboJ splicing. The presence of the plasmids with correct inserts in all clones was confirmed by Sanger sequencing (Macrogen, South Korea).

### Plasmid construction

All plasmids used to measure the autocatalytic activity of RiboJ are listed in **Table 1**. Plasmids p69A.RJ- and p69C.RJ- were generously gifted by D. Blank, University of Basel. Plasmids p69A.RJ+ and p69C.RJ+ were constructed using plasmid pMV001 (which was created beforehand), as follows: RiboJ was ordered as four 60nt single stranded oligos with each 30nt of them being homologous to either another 60nt RiboJ oligo or PCR amplified pUA66 vector (**Table S1**). These four oligos were then assembled with PCR amplified pUA66 vector using NEBuilder HiFi DNA assembly kit (New England Biolabs). The resulting pMV001 plasmid assembly mix was then used to electroporate Top10 *E. coli* cells (Invitrogen). The presence of the RiboJ was then confirmed by Sanger sequencing (Macrogen, South Korea) from colonies grown on selective LB agar plates with 50μg/ml Kanamycin.

To create inserts for p69A.RJ+ and p69C.RJ+ plasmids the *lacZ* promoter regions from p69A.RJ- and p69C.RJ- were PCR amplified. The primers used contain 17nt overhangs that are homologous to PCR amplified pMV001 vector (**Table S1**). We ligated the vector with the inserts through Gibson assembly (Gibson et al., 2009) using NEBuilder HiFi DNA assembly kit (New England Biolabs). All primers and oligos used including sequencing primers (Integrated DNA Technologies) are listed in **Table S1**. In all cases the same method for insert and vector PCR amplification from existing plasmids was used as described by (Li et al., 2011).

### Flow cytometry

Strains for flow cytometry were grown in M9 minimal media (Sigma) supplemented with MgSO_4_, CaCl_2_, 0.4% (w/v) carbon source (glucose, galactose, or lactose), and 50μg/ml Kanamycin. They were first inoculated from a glycerol stock library into a 96 well microplate using a pin replicator (Enzyscreen B.V.) and incubated at 37°C. After overnight incubation the cultures were diluted into the same fresh media with the pin replicator and incubated the same way until they reached mid-exponential phase (~4h). At that point the cells were diluted into 1x PBS with ~1% formaldehyde and kept on ice until measuring the GFP levels on the flow cytometer.

Cytometry was performed with a BD FACSCanto II and BD FACSDiva software version 6.1.3. The GFP fluorescence was measured using the 488nm laser and a 513/17nm bandpass filter.

The data from FACSDiva were exported into Flow Cytometry Standard files, and cell gating and fluorescence analysis was performed using custom R scripts (flowCore package version 2.0.1; see **Supporting Information**). Cells were gated based on their maximal kernel density of forward and side scatter values, keeping about ⅓ of all events. The modal fluorescence was calculated from gated cells as the maximal kernel density from the fluorescence signal.

### RNA isolation

RNA was isolated from four clones a day, while clones with the same genetic background were processed together on the same day. We isolated RNA from MG1655 clones twice on two different days, all other clones were isolated just once. Each strain containing one of the four plasmids (**Table 1**) was grown from a single colony overnight in 3ml of LB with 50μg/ml Kanamycin and 2mM IPTG (Isopropyl β-D-1-thiogalactopyranoside) with shaking (250 rpm) at 37°C. Because the high IPTG concentration impared growth of SC312 strain with RiboJ plasmids (i.e., p69A.RJ+ and p69C.RJ+), we grew all SC312 clones for RNA isolation in LB with 0.2mM of IPTG instead. The next day 15ml of the same fresh media in 50ml Falcon tubes was inoculated by 15μl of this overnight culture. This was incubated at the same conditions. Once the cultures reached an exponential phase (between 1.75h - 2.5h) it was placed on an ice slurry. Next we added 7.5ml of ice-cold 5% phenol in ethanol to each 15ml of culture and kept them on ice for 15min. The cultures were then spun at 7000G for 7min at 4°C, supernatant was discarded and the pellet was redispersed in 350μl of 3mg/ml Lysozyme solution (in TE buffer). After incubating for 3min, an equal volume of RNA lysis buffer was added and RNA isolated using Monarch Total RNA Miniprep Kit (New England Biolabs). Each sample was treated by DNase I twice: 1) on-column during the RNA extraction and then 2) in-tube after RNA extraction. This was done to avoid any amplification from residual gDNA during RT-qPCR. After the second treatment with DNase I the samples were column purified and concentrated using RNA Clean & Concentrator-5 kit (Zymo Research). Quality of RNA in each sample was checked on 1% agarose gel and its concentration measured on a Qubit 4 fluorometer (Invitrogen). The isolated RNA samples were then stored in −80°C freezer.

### RT-qPCR

To assess the efficiency of PCR amplification by our primers we used RNA from MG1655 strain (all four plasmids). A ten-fold serial dilution was performed on all the RNA samples up to 10^-4^ RT-qPCR was run on all the dilutions in triplicates using two different master mixes differing by the forward primer used - F1 and F2 (**Fig. 2** and **Table S1**). The total reaction volume was 20μl with 2μl of template RNA. We used SensiFAST Probe No-ROX One-Step Kit (Meridian Bioscience) and PikoReal Real-Time PCR System (Thermo Scientific) with following cycling conditions: Reverse transcription for 10min@45°C; Polymerase activation for 2min@95°C; 40 cycles of: Denaturation for 5sec@95°C and Annealing & extension for 20sec@55°C. The C_t_ values were obtained via PikoReal software version 2.2, exported into .xlsx file and converted into .csv to be further analysed using custom R scripts (see **Supporting Information**).

To assess the autocatalytic efficiency of RiboJ, the RNA from all samples was first diluted from its original concentration (~2-3μg/μl) to 20pg/μl to obtain C_t_ values between 20 and 40 and to dilute out any potential residual of gDNA (to less than one molecule per reaction). We confirmed that no amplification occurred when omitting reverse transcriptase from the master mix. Each RNA sample was then run in three or more replicates using both primer sets (**Fig. 2**) with the same conditions described above. We exported the data from the PikoReal software version 2.2 into .xlsx files, converted these into .csv, and performed all analyses using custom R scripts (see **Supporting Information**). In brief, we determined the mean C_t_ value of all the replicates for the uncut and cut RiboJ transcripts and calculated the efficiency as the ratio of the cut and uncut transcripts using the Pfaffl method (Pfaffl, 2001):

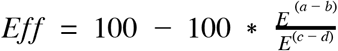

where *E* is constant mean amplification efficiency (1.95766), *a* and *b* are mean C_t_ values of transcripts without and with RiboJ, respectively, using F1 primer, and *c*and *d* are mean C_t_ values of the same transcripts using F2 primer. To obtain a measure of error in these estimates, we bootstrapped the data 10,000 times and recalculated the ratio for each bootstrap replicate.

## Acknowledgements

We would like to thank Tim Cooper for constructive input and D. Blank for providing p69A.RJ- and p69C.RJ- plasmids. This work was supported by a Marsden Grant (grant MAU1703) awarded to OKS. The funder had no role in study design, data collection and interpretation, or the decision to submit the work for publication.

## Author contributions

MV and OKS designed the study, developed the experimental methods, interpreted the data and wrote the manuscript with input from BRM. MV analyzed the data. MV and BRM performed the experiments.

## Data availability

The original data files and scripts with data analysis that support the findings of this study are available in the Supporting Information of this study.

## Supporting information

All scripts and data files can be found at https://github.com/marketavlkova/RiboJPaperSupp

**Table S1:**
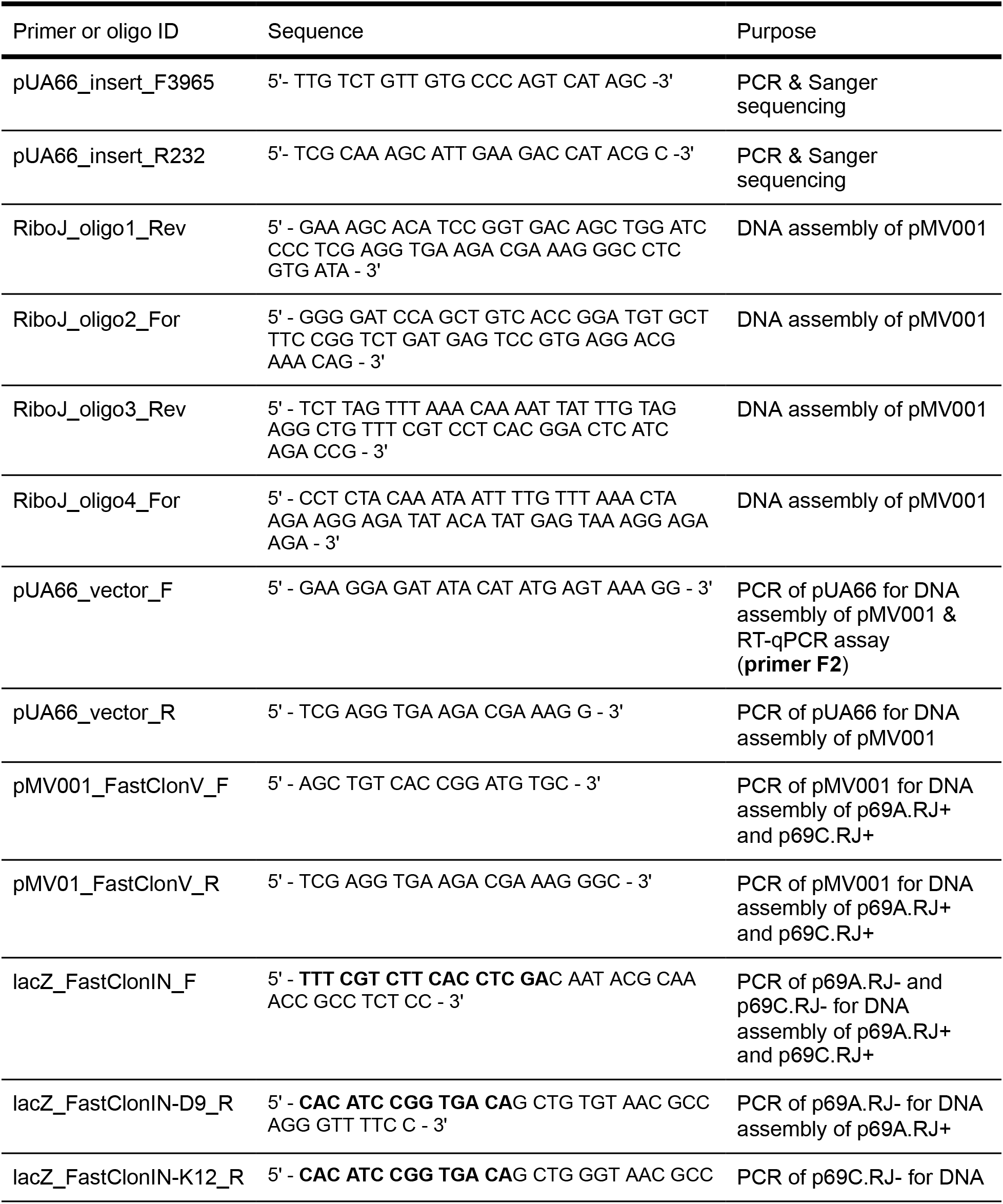
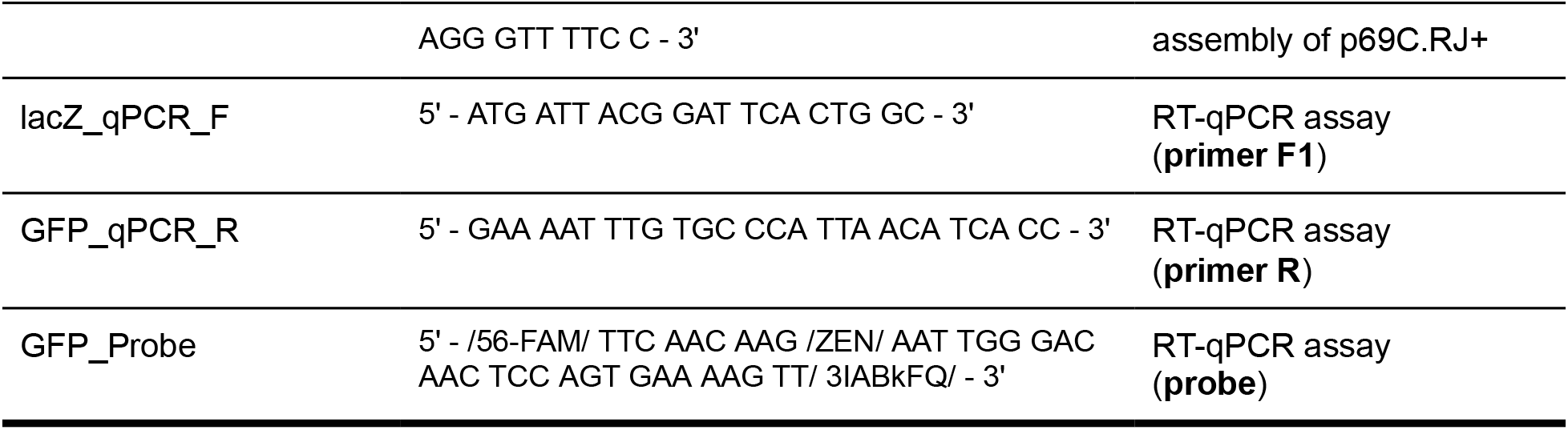
Primers and oligos used in this work. Bold sequences = regions homologous to PCR amplified pMV001 vector. Primers and the probe used for RT-qPCR assay are highlighted in bold with their simplified names that were used in the main text and Figure 2 (Purpose column).

**Figure S1:**
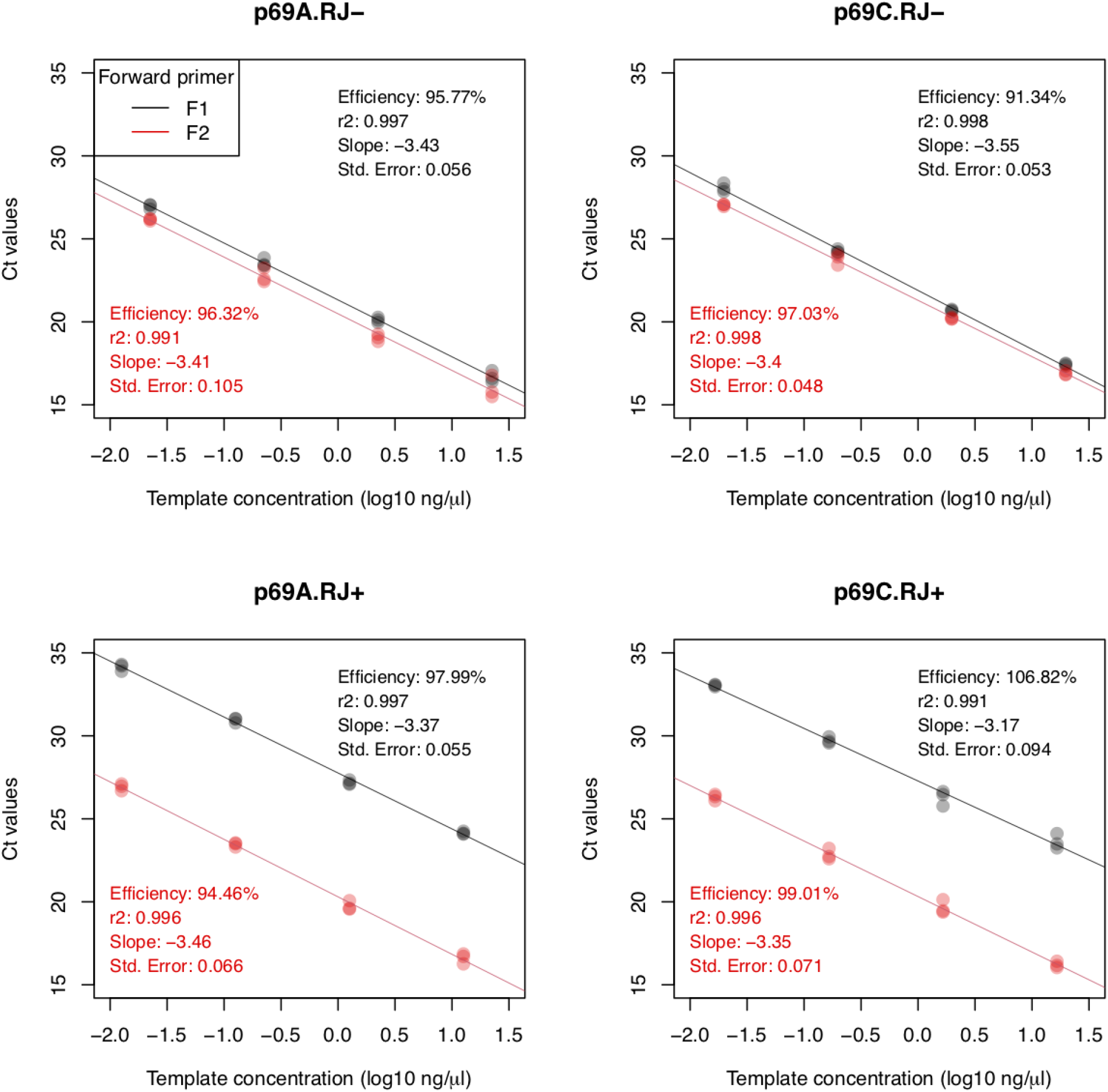
Amplification efficiency of all primer-template combinations. Each point indicates the C_t_ value for one technical replicate at different template concentrations, with each panel indicating one template and each color (red or black) indicating one primer combination. The lines show linear regressions, calculated using all data points for a given primer-template combination. For the templates without RiboJ (top panels), both primer pairs result in nearly equal C_t_ values. For the templates with RiboJ (bottom panels), the F1 primer pair has consistently larger C_t_ values, as expected. To calculate the catalytic activity of RiboJ, we used the mean amplification efficiency across all primer-template combinations, 95.8%.

